# Multiple splitting of droplets using multi-furcating microfluidic channels

**DOI:** 10.1101/513812

**Authors:** Zida Li, Luoquan Li, Meixiang Liao, Liqun He, Ping Wu

## Abstract

Removing volumes from droplets is a challenging but critical step in many droplet-based applications. Geometry-mediated droplet splitting has the potential to reliably divide droplets and thus facilitate the implementation of this step. In this paper, we report the design of multi-furcating microfluidic channels for efficient droplet splitting. We studied the splitting regimes as the size of the mother droplets varied and investigated the effect of channel lengths. We further examined the droplet breakup mechanisms. This study could potentially be applied in washing steps in droplet-based biological assays or assays that require aliquot.

## Introduction

Homogeneous nanoliter/picoliter droplets have great applications in the fields such as food industry^1^, cosmetics^2^, single cell RNA/DNA sequencing^3^, droplet digital PCR^4, 5^, nanoparticle synthesis^6^, drug delivery^7^, and drug discovery^8^. These droplets offered a platform for precise control over reagent quantity, which are critical for reaction reproducibility and reliability. Additionally, cells and objects can be encapsulated in the droplets, providing a mean to manipulate single cells^9, 10^.

Traditionally, droplets are prepared by mechanical agitation. Despite its wide applications, this method suffered from poor droplet monodispersity, leading to compromised process controllability. Leveraging the strict laminar flow and Plateau-Rayleigh instability, microfluidics has been proved as a powerful tool for the generation of highly monodispersed droplets, typically with a coefficient of variation in droplet size between 1% and 3%^11^. In addition, the throughput of droplet generation is high: hundreds to thousands of droplets can be generated each second, depending on the flow conditions. Many chemical and biological applications have been implemented in droplet-based microfluidics, such as drug screening, enzyme revolution, particle synthesis, material synthesis, and single cell sequencing^9, 10, 12, 13^.

In a typical application of droplet-based microfluidics, chemicals or objects are encapsulated within individual droplets during droplet formation. Many downstream techniques have been developed to achieve specific droplet manipulation. For example, droplet sorting techniques was developed for encapsulation enrichment^14^, droplet merging for large volume reagent exposure or object contact^15^, and pico-injection for small volume reagent addition^16^. These techniques served as powerful tools for downstream droplet manipulation. Nevertheless, most of these techniques focused on volume addition. Volume removal, on the other hand, is equally important in many application scenarios. For example, many biological assays, such as immunoassays, incorporate one or more washing steps, requiring both reagent addition and removal. When performed in droplets, these assays can be implemented by encapsulating microbeads, which serve as attaching substrates. By adding and removing reagents from the droplets, washing steps can be implemented. Therefore, efficient volume removal techniques are critical for these applications.

Many works have been reported to achieve volume removal from droplets by actively splitting droplets based on methods such as dielectric^17^, pneumatic^18^, magnetic^19^, thermocapillary^20^, and acoustic forces^21^. However, these methods were featured with extra physical fields, thus limiting their applications in practice. With carefully designed T-junctions in microfluidic devices, passive droplets splitting mediated by geometry and the resultant hydraulic resistance were reported^22, 23^. This method produced daughter droplets with designed sizes and good reproducibility. Nevertheless, such binary splitting has limited efficiency for applications where multiple aliquoting and thus multiple splitting is necessary. To address this problem, herein we report a simple yet robust droplet splitting mechanism which offers tunable amount and splitting ratio among the daughter droplets. In particular, we adopted multi-furcating channels to break up the mother droplets and direct the daughter droplets. When the mother droplets came to the furcating region, hydraulic pressure deformed them into narrow rod-like shapes, facilitating the breakup into daughter droplets. We investigated the effect of mother droplet sizes; we found that as the droplet size increasing, the droplet splitting fell in different regimes, including no-splitting, non-plug splitting, and plug splitting regime. We further investigated the effect of flow resistances on splitting results. Our results showed that by adjusting channel lengths, the splitting results could be tuned. We further investigated the splitting mechanisms. Our observation showed that the splitting was principally resulted from the mother droplet deformation induced by the colliding on the channel edge. However, when the mother droplet was extensively deformed into a rod with very small diameter, spontaneously breakup would happen, very likely due to Plateau-Rayleigh instability. This droplet splitting method holds great potential as a versatile tool in droplet-based biological assays which require washing or sample splitting.

## Materials and Methods

Microfluidic devices were fabricated by standard photo lithography and polydimethylsiloxane (PDMS) replica molding. Briefly, SU8 (2030, MicroChem Corp.) was spin-coated on silicon wafer, typically with a thickness of 5 μm, before it was exposed by UV and developed. The fabricated silicon wafer with SU8 structure was then silanized before it was used for PDMS (Sylgard 184, Dow Chemical Company; 10:1 base-to-hardener ratio) replica molding. The cured PDMS was then cut and peeled off from the silicon wafer. Holes with diameters of 1 mm were punched using biopsy punches. The PDMS slab was then bonded with glass slide after oxygen plasma treatment (PDC-32G-2, Harrick Plasma) for 1 minute and baked for 5 minutes.

Chemical reagents were loaded in 1 mL glass syringes (Hamilton Company) and infused into the microfluidic chips by syringe pumps (NE-1200, New Era Pump Systems Inc.) via polytetrafluoroethylene (PTFE) tubing (1522, IDEX Health & Science LLC). An inverted microscope (IX73, Olympus Corp.) coupled with high speed camera was used to observe and image the experiments.

Mineral oil (M3516-1L, Sigma Aldrich) used as the continuous phase, with 0.05% Triton X-100 (T9284, Sigma Aldrich) and 2% ABIL EM90 supplemented as the surfactant. DI water was used as the disperse phase.

Diameters of the generated droplets were measured using ImageJ (National Institute of Health, USA).

## Results

### Overview of the splitting setup and demonstration of the splitting performance

Mother droplets with uniform size were generated with T-junctions, as shown in Figure S1, typically with a coefficient of variation of less than 5% in droplet diameter^24^. A cross junction was designed at the downstream to break up the mother droplets and direct the daughter droplets towards designated outlets (Figure 1a). Upon arriving at the cross junction, due to pressure drop and shear stress, the droplet was deformed into a rod shape with significantly reduced diameter. The area around the notch between adjacent channels has elevated pressure, which squeezes the droplet and leads to locally thinned rod diameter. Eventually, the droplet breaks up at the loci of the droplet close to the notches.

**Figure 1.**
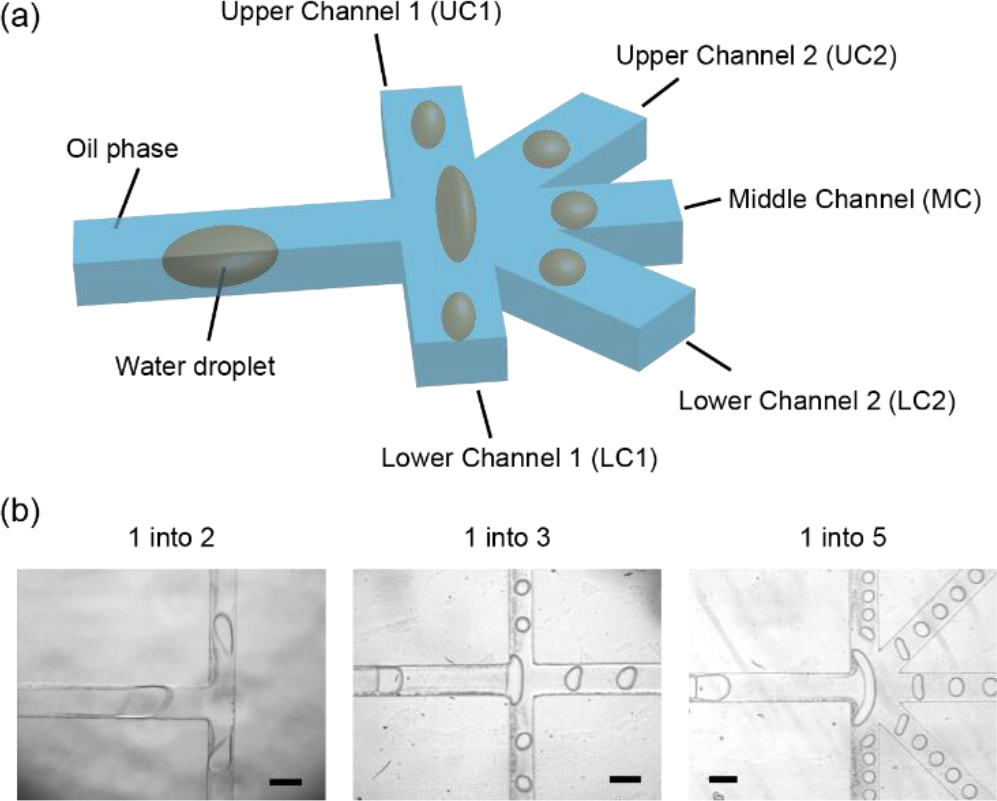
(a) Schematic showing the droplet splitting setup. A droplet coming downstream to the junction splits into a few daughter droplets. (b) Representative images showing droplets split into two, three, and five daughter droplets, as indicated. Scale bar, 200 μm. Flow rates of the continuous phase and disperse phase were 6 and 1, 10 and 1, and 2 and 0.5 μL/min, respectively.

By adjusting the channel design, splitting of droplets into two, three, and five daughter droplets were achieved, respectively (Figure 1b). Daughter droplets showed symmetry along the axis of the main channel: droplets moving towards north and south displayed similar droplet sizes, and these moving towards northeast and southeast displayed similar sizes. This observation agreed with the symmetry of the channel designs. Among each branch channel, the droplets showed high uniformity, with coefficients of variation in the droplet diameters typically less than 5%.

The splitting results largely depend on the pressure distribution of the flow field, which is the coupled result of a few parameters including flow rates, viscosities of the fluids, interfacial tension, size of the mother droplet, cross junction geometry, flow resistance of the branch channels, etc. Size of the mother droplet, in particular, is dependent on the flow rates, fluid viscosities, and interfacial tension. To simplify the study, we focused the study on the effect of size of the mother droplets and flow resistance of the branch channels.

### The effect of the size of mother droplets on the resultant daughter droplets

The size of the mother droplet affects its shape at the junction, critically affecting the splitting results. The size of the mother droplets can be adjusted by tuning the flow rates of the continuous phase (oil) and disperse phase (water)^24–26^. We investigated the effect of mother droplet size on splitting in channels with one-into-five star-shape junctions by varying the mother droplet size from 50 to 350 μm. Results showed that when the mother droplet was small, typically with a diameter below 70 μm, it was slightly deformed at the junction but entered only the Middle Channel with no splitting, as illustrated in Figure 2. We named this regime no-splitting regime.

**Figure 2.**
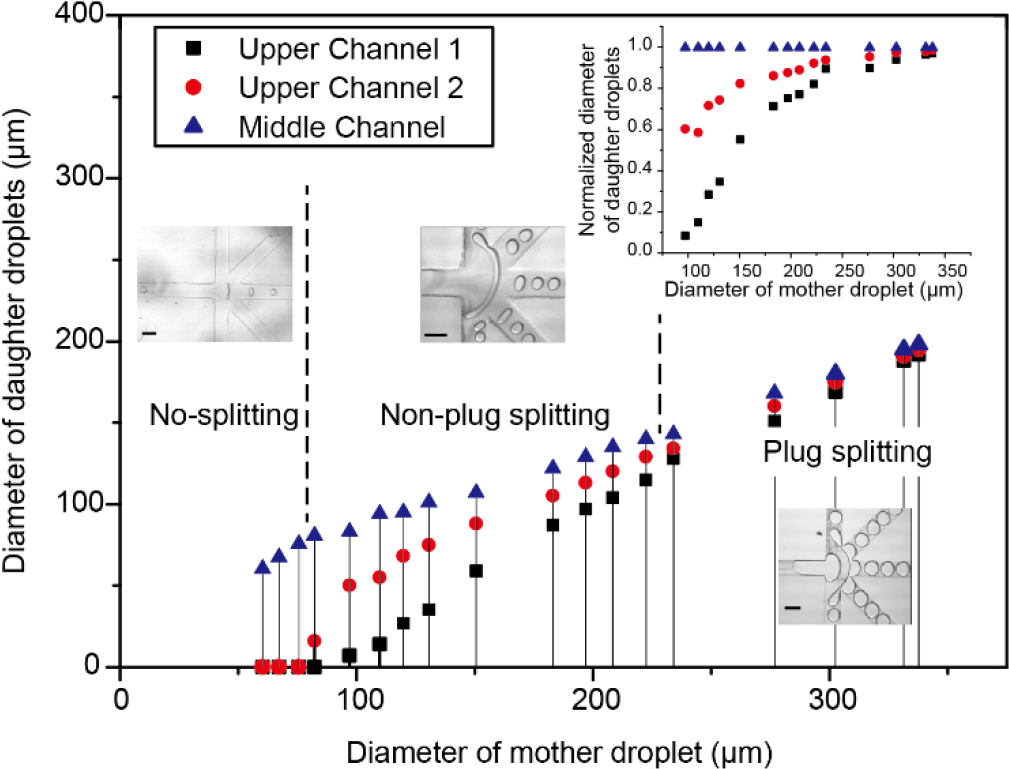
Droplet splitting fell into three regimes, namely non-splitting, non-plug splitting, and plug splitting, as diameter of the mother droplet increases. In plug splitting regime, daughter droplets showed similar sizes. Channel names were designated in Figure 1a. Scale bars, 200 μm. The flow rate of the continuous phase was fixed at 2 μL/min while that of the disperse phase varied from 0.1 to 1 μL/min. Inset: Normalized diameters of daughter droplets, which were the ratios of droplet diameters to those of Middle Channel, plotted as a function of the diameters of mother droplets.

As the droplet size increased, the two channels adjacent to the Middle Channel, namely Upper Channel 2 and Lower Channel 2, started to receive daughter droplets. When the droplet diameter reached about 100 μm, the mother droplet split into five daughter droplets, directed into five branches, respectively. Droplets in mirrored channels, namely Upper Channel 1 and Lower Channel 1, and Upper Channel 2 and Lower Channel 2, typically had similar droplet sizes. The diameters of the daughter droplets in MC were always the largest among those of the five channels. Daughter droplets in Channels 1 and 2 had much smaller diameters than those in MC. However, this difference was narrowed as the diameter of the mother droplets increased. When the diameters of the mother droplet reached about 230 μm, the daughter droplets had almost same diameters, as shown in the inset of Figure 2. We noticed that at this stage, mother droplets obstructed the continuous phase and formed a plug flow, and the daughter droplets also covered up the whole cross section of the branch channels, forming plug flows as well. As such, we named the former regime as non-plug splitting regime and the latter regime as plug splitting regime, as indicated in Figure 2.

The observation of equal-sized daughter droplets in plug splitting regime could be explained by the linear dependence of daughter droplet size on the flow rates of branch channels in plug flows. The mother droplet filled up the channel cross section and segmented the continuous phase. As it approached the junction, the distribution of its volume in each branch channel would be proportional to the flow rate of each branch channel. In a continuous flow, the flow rate in branch channels are largely decided by the flow resistance, including local resistance at the junction towards each branch channel and Darcy friction in each branch channel. As the mother droplet would fill up the whole junction and each branch channel had equal length thus equal Darcy friction, mother droplets evenly split into five daughter droplets and flowed into each of the five branch channels. Similar results were also observed in one-into-three junction structures (Figure S2).

### Effect of channel lengths on splitting ratio in no-plug splitting regime

Though in plug splitting regime the splitting ratio is relatively predictable: mother droplets would split evenly into each branch channel, in non-plug splitting regime, the splitting ratio is more diverse. Since the splitting process depended on the flow field and thus flow resistances of the branch channels, we asked whether the sizes of the daughter droplets could be controlled by varying the lengths of the branch channels. Using one-into-five junction designs, we varied the length of the side channels. Decreasing the lengths of the side channels decreased their flow resistance, resulting in smaller pressure drop between the main channel and side branches. Consequently, the daughter droplets split into these channels would have large sizes.

Indeed, when UC1/LC1had a length of 2 mm, while UC2/LC2 and MC had lengths of 5 mm, the diameters of the daughter droplets in UC1/LC1 were significantly larger than those in UC2/LC2 and MC, as shown in Figure 3a. Similarly, when UC2/LC2 were 2 mm long, while keeping UC1/LC1 and MC at 5 mm long, the diameters of the daughter droplets in the UC2/LC2 were much larger than UC1/LC1 and MC, as shown in Figure 3b. Furthermore, by increasing the resistance of MC and decreasing that of UC1/LC1, we were able to avoid daughter droplets entering MC. For example, when the lengths of UC1/LC1, UC2/LC2, and MC were 2 mm, 5 mm, and 7mm, respectively, we observed larger daughter droplets in UC1/LC1 but daughter droplets were absent in MC.

**Figure 3.**
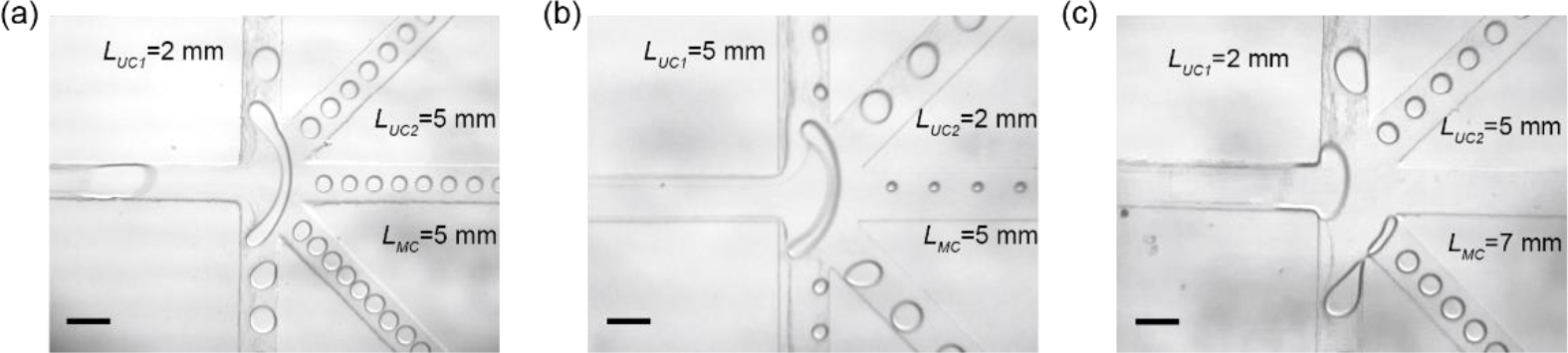
Effect of channel length on resultant child droplets. Lengths of the branches decided the flow resistances, leading differed flow rates and thus droplet size. Channel lengths were indicated in the figure, with upper and lower half being symmetric. Flow rates of the continuous and disperse phase were 2 and 0.5 μL/min, respectively. Scale bars, 200 μm.

These results showed that by adjusting the lengths of branch channels, we could tune the splitting ratio. Since droplet microfluidics is generally not plug flow and droplet splitting usually lies in non-plug splitting regime, this observation could have profound applications. We investigated whether this device could achieve on-demand droplet splitting with carefully designed channel length. Specifically, we studied the parameters that could generate equal-sized droplets in the five channels by adjusting the lengths of UC1/LC1 and MC while keeping UC2/LC2 at 5 mm. As we set the lengths of UC1/LC1, UC2/LC2, and MC as 4.1, 5, and 5.7 mm, respectively, the diameters of the resultant daughter droplets were very close (Figure 4). The measured diameters of the droplets in UC1/LC1, UC2/LC2, and MC were 108.6 ± 3.1, 104.6 ± 2.6, and 108.8 ± 4.3 mm, respectively. These results showed that by careful design, we could obtain desirable droplet splitting ratios on-demand.

**Figure 4.**
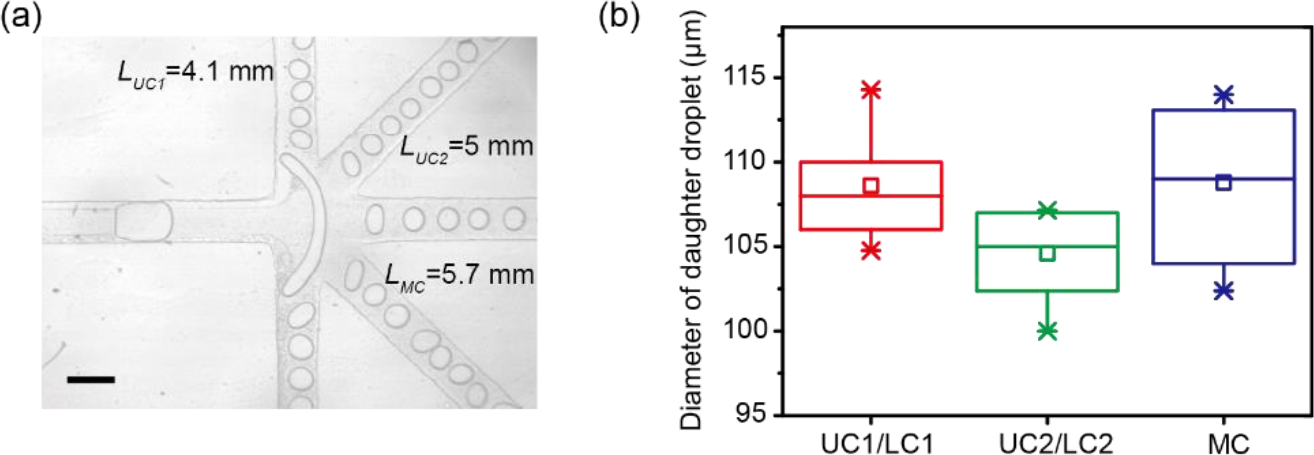
Generation of droplets with same sizes in all channels by tuning the channel lengths. (a) Microscopic images showing equal-sized droplets generated in each branch channels. Scale bar, 200 μm. (b) Bar plot of the diameter of the daughter in each branch channels.

### Breakup mechanisms

Mother droplets were deformed into rod shape when it passed through the junction due to the pressure drop towards branch channels across the droplets, as illustrated in Figure 5a. The deformed droplets were subject to shear stress and surface tension, which were balanced to maintain the integrity of the mother droplet. In the meanwhile, the droplet surface was subject to Plateau-Rayleigh instability, which disturbed the surface with minute sinusoidal waves. Plateau-Rayleigh instability stated that a liquid cylindrical jet in infinite space is unstable and eventually break up into drops; the characteristic breakup time is proportional to *R*_*0*_^3/2^, where *R*_*0*_ is the diameter of the original jet [Citation John Bush Surface Tension Module Lecture 5]. In a confined space, such as this case in microfluidics, though it is very difficult to obtain analytical results, one would expect that the scaling law between breakup time and jet diameter applies at least qualitatively. Thus, at large rod diameters, instability induced surface disturbance did not develop fast enough, resulting in collision of the mother droplets on the channel edges (Figure 5b). Once the elongated droplet collided with the any corner of the junctions, a neck was formed and the collapse could easily occur. The breakup mechanism was very similar as those of using T or Y junction for droplet generation. The viscous stress was the main force promoting the droplet breakup. It was proved that the occupation of dispersed liquid in T junction led to the reverse flow around the surface and it determined when a neck collapses rapidly to form a droplet^27^. This breakup mechanism had a feature that the breakup always occurred near to the channel corner. Thanks to the laminar flow and high flow reproducibility in microfluidics, the resultant daughter droplets had good consistency in each branch channels.

**Figure 5.**
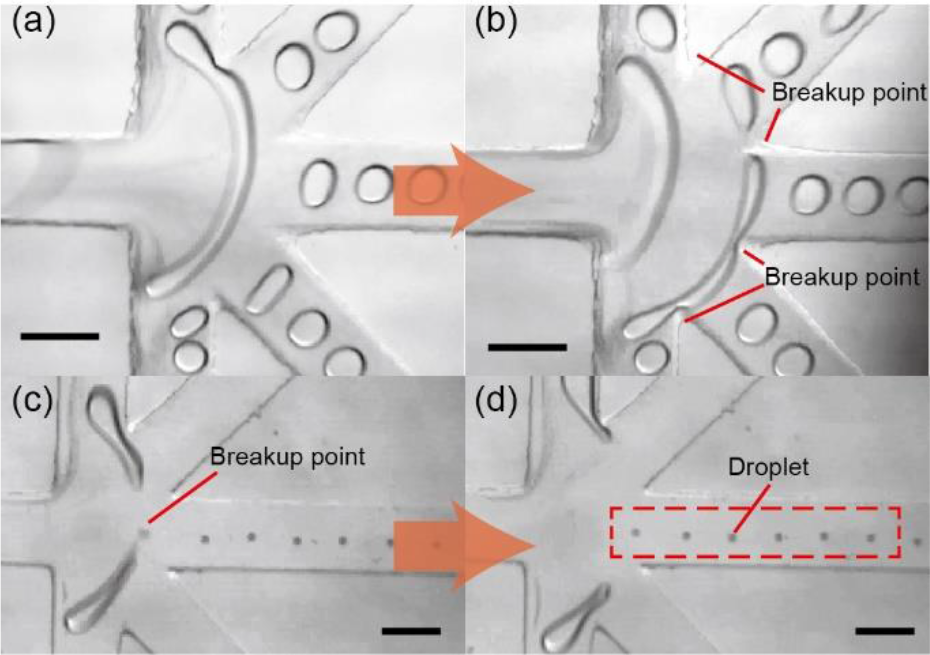
The two observed breakup mechanisms. (a & b) Mother droplet was elongated into a rod-like shape and collide with channel edge, breaking up into daughter droplets. Flow rates of the continuous and disperse phases were 1 and 0.2 μL/min, respectively. (c & d) When the diameter of the deformed rod-shaped mother droplet was small enough, droplets would be generated at place even without colliding with the channel edges, due to Plateau-Rayleigh instability. Flow rates of the continuous and disperse phase were 2 and 0.15 μL/min, respectively. Scale bars, 200 μm.

The other breakup mechanism observed was much more influenced by extension stress. As described above, the mother droplet could become very thin before collided with the channel edges if the deformation happened fast (Figure 5c). With small rod diameters, Plateau-Rayleigh instability became more predominant, leading to breakup ahead of collision. This breakup mechanism was commonly observed in one-to-two splitting using T junction and droplet generation in co-flows, and the breakup point was determined by thinnest location instead of solely by the channel edges. In this mechanism, the breakup location was less reproducible and daughter droplets with small size were generated, as shown in Figure 5d. Here, remarkably shortening the length of UC1/LC1 could induce high flow rates into these channels and greater extensional stress and thus high deformation, resulting in this breakup mechanism.

## Conclusion

In this paper, we reported the observation of droplet splitting in microfluidic channels. In particular, we fabricated furcating channels and split each mother droplet into five daughter droplets in a consistent manner. We investigated the effect of the sizes of mother droplets and found that as the size of the mother droplet increased, the splitting fell into three regimes, namely non-splitting, non-plug splitting, and plug splitting regime. We further studied the effect of flow resistance in the branch channels, particularly the lengths of branch channels, on the droplet splitting. We found the channel lengths that could lead to equal-sized daughter droplets formation. The breakup mechanism was very similar to that of T-junction for droplet generation if the breakup occurred at the solid corner. However, Plateau-Rayleigh instability played a more significant role when the droplet was intensively stretched and thinned due to highly unbalanced flow resistance in the branch channels. This study provided a proof-of-concept method for volume splitting in droplet-based applications where aliquot or washing is difficult to implement.

## Acknowledgements

This work is supported by National Natural Science Foundation of China (NSFC 31500694 and NSFC 31670866).

