## Supplementary Information for "Multiple splitting of droplets using multi-furcating microfluidic channels"

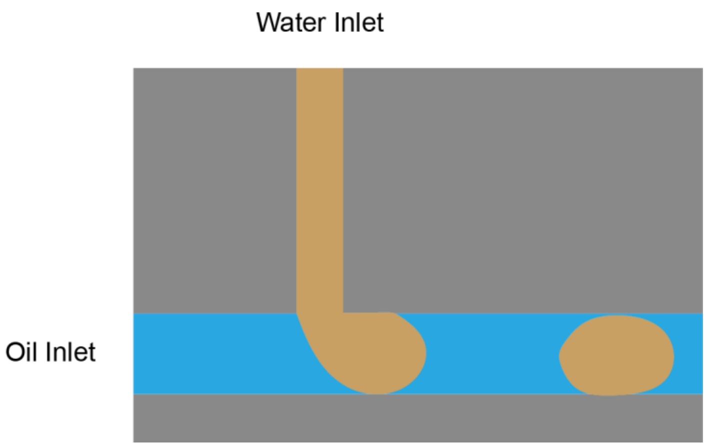


Figure S1. Generation of mother droplets using T-junctions.

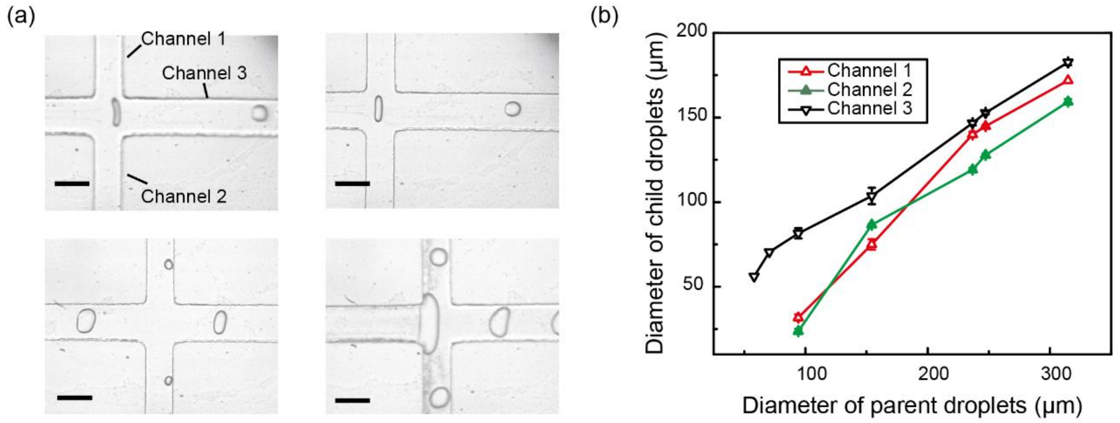


Figure S2. The effect of mother droplet size on the resultant daughter droplets in one-to-three junctions. (a) The amount of daughter droplets increased from one to three, as the size of mother droplets increased. Scale bar, 200 μm. (b) The diameters of daughter droplets plotted as a function of the diameter of mother droplets. Channel names are designated in Figure S2a.
